# Bacterial growth-promoting properties of pooled urine and individual urines from post-menopausal women with or without a urinary tract infection can vary substantially

**DOI:** 10.1101/2022.05.26.493673

**Authors:** Jacob Hogins, Ethan Fan, Zheyar Seyan, Sam Kusin, Alana L. Christie, Philippe E. Zimmern, Larry Reitzer

**Author notes:** Address correspondence to Larry Reitzer.

## Abstract

Urinary tract infections (UTIs), mostly caused by uropathogenic *E. coli* (UPEC), are common bacterial infections that affect a majority of women and often recur. Genomic and transcriptomic analyses have not identified a common set of virulence genes which has suggested conserved virulence functions instead of virulence genes, multiple virulence mechanisms, and complex host-pathogen interactions. One aspect of the host-pathogen interaction is rapid UPEC growth in urine in vivo. When bacteria are grown in pooled urine, an averaged urine is assumed to diminish individual variation. We grew a non-pathogenic and pathogenic *E. coli* strains in urine from individuals who never had a UTI, had a UTI history but no current infection, and had a UTI history with a current infection. Bacterial growth showed large variations in individual urines and pooled urine supported significantly more growth than predicted for never and history groups but not the current group. UPEC strains, but not the non-pathogenic strain, were resistant to urinary inhibitory factors e.g., antimicrobial peptides based on an indirect inoculation-density effect assay. Total nutrient content tended to be higher in current group urine than never and history group urine. We propose that pooling optimizes a nutrient mixture in never and history group urines, which are often studied, whereas urine from current group individuals appear to have a more optimal nutrient mixture. We conclude that pooled urine is not “an average urine”, and that the best comparisons of results between labs using pooled urine would also include results with a standardized synthetic urine.

## INTRODUCTION

Urinary tract infections (UTIs) are the most common bacterial infections in humans. 50% of women will have a UTI by age 35 (1), and 20% of women in the 18-24 age group will have a UTI annually (2, 3). After an initial infection, the risk of recurrence is 25-30% in the ensuing 6 months (4). Recurrent UTIs (RUTIs) are more common in postmenopausal women, can last for years, and become harder to control, in part, because of antibiotic recalcitrance (allergy and/or resistance) (5, 6). Uropathogenic *E. coli* (UPEC) causes 80-90% of community-acquired UTIs and most RUTIs (7, 8).

UTI progression includes bacterial entry into the bladder, rapid growth in urine, attachment to and entry into uroepithelial cells, rapid replication to form intracellular bacterial communities, and release back into the bladder where the bacteria can reenter uroepithelial cells or penetrate into deeper bladder layers (9). Several genomic analyses have attempted to identify putative urovirulence factors required for these processes, but a recent and thorough genomic analysis failed to identify a common set of virulence factor genes in UPEC (10). This failure led to the core gene expression hypothesis: some genes that are present in all *E. coli* strains are more highly expressed in UPEC strains. However, the follow-up transcriptomic analysis also failed to identify specific genes whose expression correlated with symptom severity in a mouse model (10). The apparent failure of genomic and transcriptomic analyses to identify virulence-specific genes led Hultgren and colleagues to propose that UTIs are a complex interplay between bacterial and host properties, and that the shared features of UPEC strains are not specific genes or their expression but conserved functions, such as carbohydrate metabolism and motility (10). Mobley and colleagues have proposed extremely rapid UPEC growth in urine and high expression of genes associated with rapid growth as conserved functions (11, 12).

One aspect of the host-pathogen interaction is bacterial growth in urine, which could conceivably depend on simultaneous utilization of multiple energy-generating nutrients, such as low levels of amino acids and carbohydrates (13). Urine also contains a variety of defense mechanisms that impair disease progression (14). Studies that characterize bacterial growth in urine usually involve pooled urine which is assumed to eliminate variations between individuals and allow reproducibility between labs. These assumptions imply limited variation between individuals and that an average urine exists. These assumptions are untested and could be wrong if large variations exist for nutrient content and host defense mechanisms between individuals.

To assess variability between urine samples and, therefore, to determine whether results using pooled urine are reproducible, we compared bacterial growth in a pooled urine to the average growth in individual urine samples. Our results suggest that pooled urine is not necessarily an averaged urine, and the basis for this discrepancy is likely to be variations in nutrient content. These results suggest that reproducibility is not guaranteed for labs that study or use bacterial growth in pooled urine.

## RESULTS

### The properties of pooled urine and individual urines can vary substantially

Aspects of pathogenicity have often been analyzed after bacterial growth in pooled urine and Table 1 summarizes pooled urine characteristics from eight studies. The pooled urine was from 2-10 individuals (average 5.6). When details of the study participants were provided, which was not always the case, the participants were described as healthy which often means not having had a UTI within the past few months, and in three studies, urine from both men and women were pooled.

**Table 1.** Compilation of pooled urine studies

| Description of pool | Start<sup>a</sup> | Doubling time<br>(minutes) | Purpose of experiment | Ref |
| --- | --- | --- | --- | --- |
| 3-5 women, ages 20-40, no recent UTI | Not stated | Not given | Microarray transcriptomics | (29) |
| ≥ 3 men and women, no recent UTI | 0.05 | 50-100 | Comparative growth of bacterial strains | (30) |
| 8-10 men and women | ~0.01 | 40 | Comparative proteomics | (31) |
| 8-10 men and women | ~0.15 | Not given | Comparative proteomics | (32) |
| 5 healthy donors | ~0.004 | Not given | Microarray transcriptomics | (33) |
| 3-5 women, no recent UTI | 0.05 | Not given | Microarray transcriptomics | (34) |
| ≥ 3 women, no recent UTI | 0.01 | 50 | Comparative growth of bacterial strains | (35) |
| ≥ 2 healthy volunteers | unknown | Not given | Analysis of pili synthesis | (36) |

a. Numbers refer to starting OD600. An estimate, indicated with “~”, is given based on results presented in the paper

Based on these studies, we tested whether pooled urine from five women supported the same growth as the average growth from individual samples. Mid-stream urine samples were collected from post-menopausal women in a “never” group in which women self-reported never having a UTI, a “history” group in which women had a history of UTIs but no current infection (asymptomatic and negative urinalysis), and a “current” group in which women had a history of UTIs and an active symptomatic UTI. We grew three bacterial strains in each urine sample: W3110, a nonpathogenic lab strain; UTI89, a highly passaged model UPEC strain (15); and LRPF007, a recently isolated UPEC strain from a patient with RUTIs. We analyzed the ΔOD600 as a measure of growth instead of doubling time because few cultures had a constant exponential growth rate for more than one generation even with 30-minute sampling intervals. A possible reason for the lack of extended exponential growth is because bacteria were grown in microtiter dishes, which is required for multiple assays, and such growth was undoubtedly microaerobic; however, this may be more physiologically relevant since the bladder is a microaerobic environment as well (16, 17). In addition, the extent of growth allows an assessment of the inhibitory factors (e.g., antimicrobial peptides, etc) as described in the next section.

We examined the effect of pooling by determining the growth ratio in pooled urine versus the average growth in individual urines. We define this ratio as the PI (pooled versus individual) ratio, with a ratio of 1.0 indicating the absence of a pooling effect. The PI ratio often deviated from 1.0 (Figure 1). The greatest ratios were from UPEC growth in never and history group urines, for which the PI ratios ranged from 1.74 to 3.05 and the differences were statistically significant (p < 0.05) (Table 2). UPEC growth in pooled urine could be greater than growth in any single urine sample (Figure 2). In contrast, the PI ratios were ≤ 1.1 for UPEC growth in current group urine and generally for W3110 growth. To understand the basis for the growth synergism, we examined the two general types of factors that affect bacterial growth in urine: growth inhibitory factors and total nutrient content.

**Figure 1.**
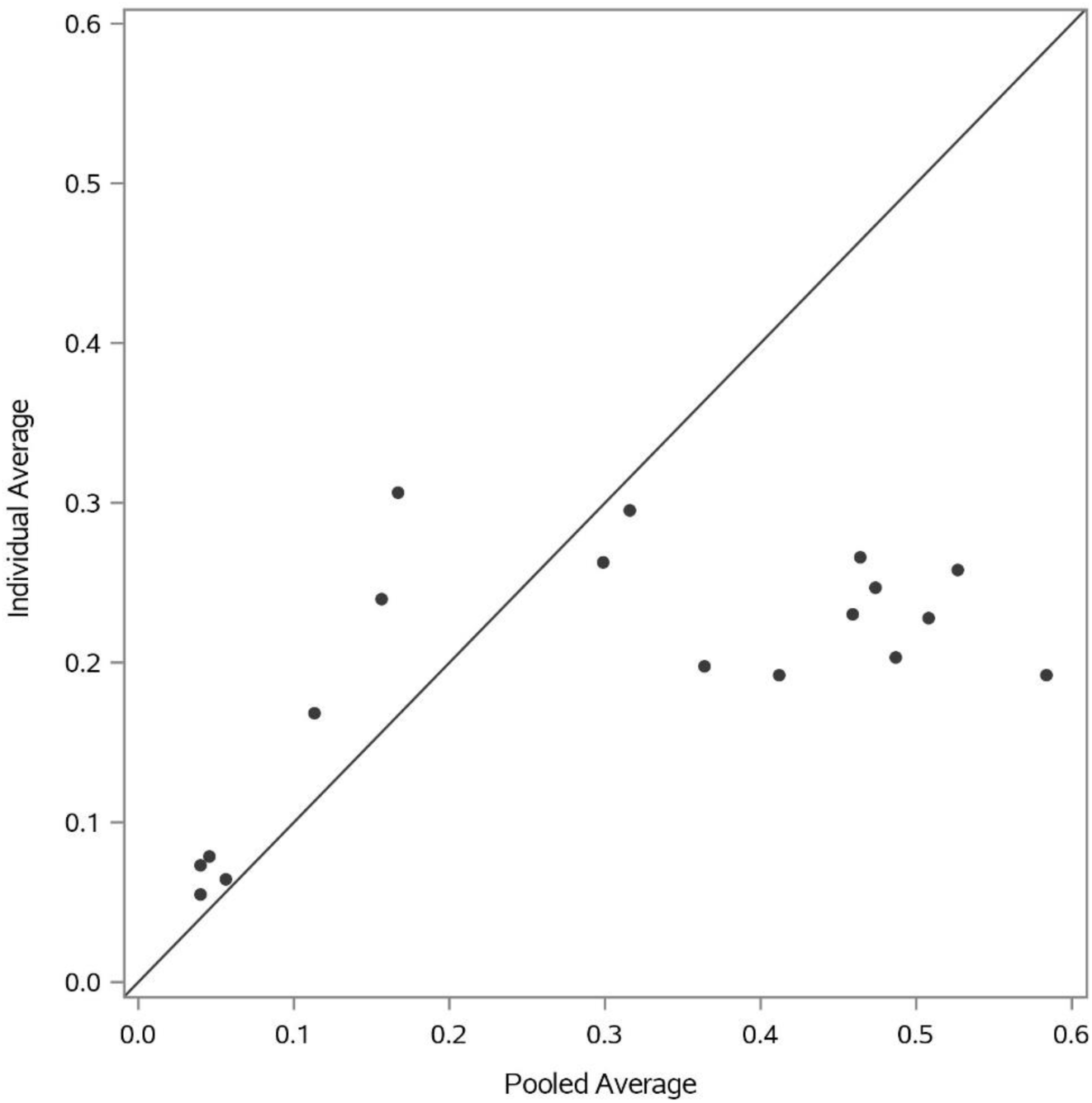
Growth in pooled urine versus the average growth from individual urines. The numbers refer to the ΔOD600. The pooled urine was from five individuals and the average is from the same five individuals. The diagonal line is expected if the ratio is one, i.e., growth in pooled urine is the same as the average of the individual urines.

**Table 2.** Comparative growth in pooled urine versus the average of individuals. Growth in each individual urine is the average of a triplicate growth experiment. Each group has 5 samples, and the 5 triplicate averages were then averaged to provide the average growth in the individual urine samples. Growth in the pooled urine is performed in triplicate, and the 3 values are used for statistical comparisons. F-tests were used to analyze if variance was unequal between the individual versus pooled urine samples. If the F-test was significant, the heteroscedastic *t*-test was used to test for differences in the means; if the F-test was not significant, the homoscedastic *t*-test was used.

|  | W3110 |  | UTI89 |  | LRPF007 |  |
| --- | --- | --- | --- | --- | --- | --- |
| Inoculation | low | high | low | high | low | high |
| Current | 0.88 (0.72) | 0.67 (0.42) | 0.65 (0.39) | 1.07 (0.85) | 1.14 (0.74) | 0.54 (0.25) |
| History | 0.55 (0.24) | 0.58 (0.14) | 1.74 (0.03) | 1.99 (0.008) | 1.92 (0.0007) | 2.23 (0.005) |
| Never | 0.72 (0.18) | 1.83 (0.015) | 2.39 (0.005) | 2.15 (0.003) | 3.05 (0.003) | 2.04 (0.005) |

**Figure 2.**
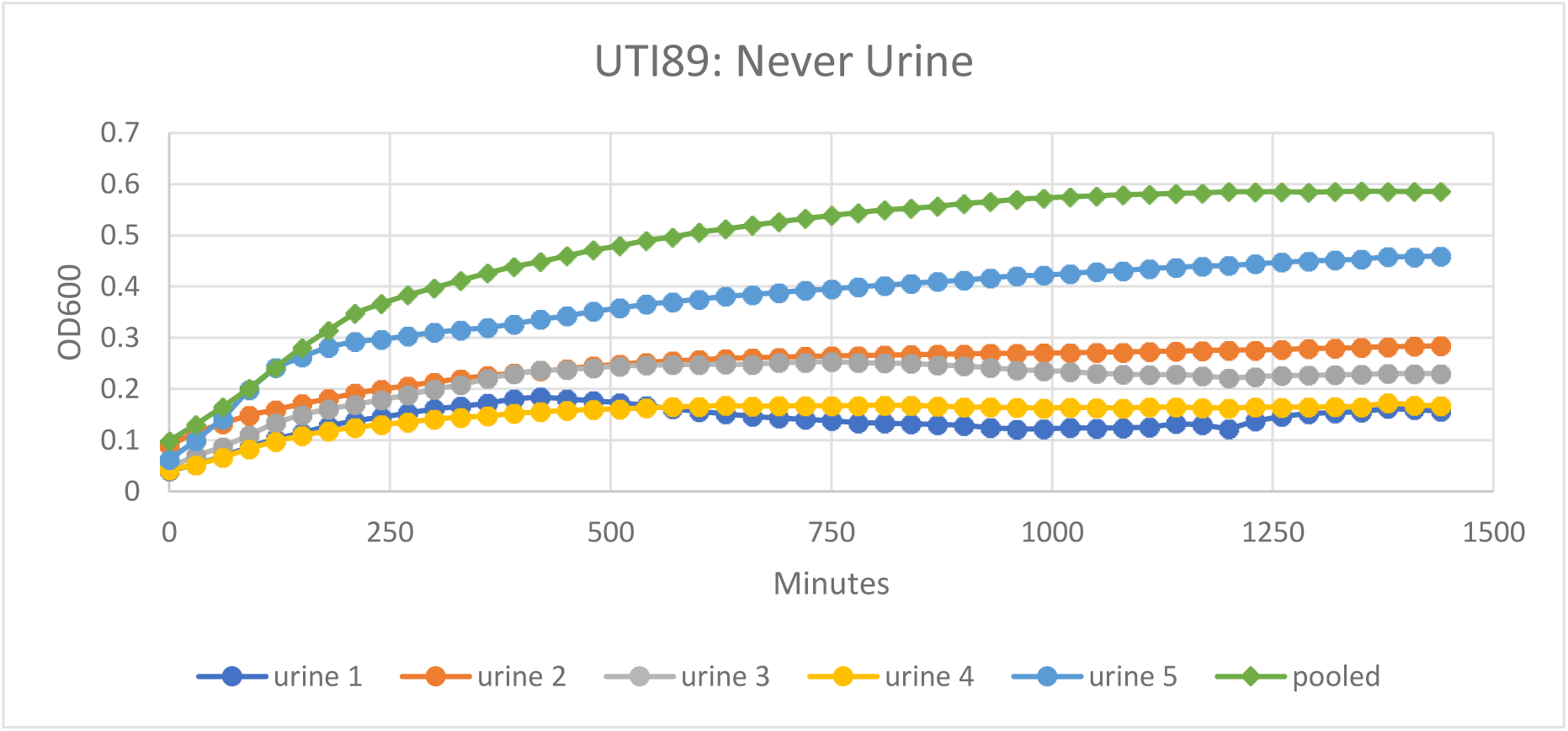
Growth in pooled urine can be greater than any single individual urine. The example is for UTI89 growth in never group urine.

### Growth inhibitory factors, the inoculation-density effect, and UPEC resistance

A characteristic feature of growth inhibitory factors, such as antimicrobial peptides and antibiotics, is the so-called inoculation density effect — the initial inoculation density determines the final cell density — which was first observed in 1946 (18). All explanations of this effect involve overcoming the negative consequence of an inhibitory factor (IF) (18).

To confirm that the inoculation effect can result from IFs, we examined whether the presence of the antimicrobial peptide LL-37 results in an inoculation effect in two media (Figure 3). For both W3110 and UTI89, the inoculation effect was greatest at 4 µM LL-37 but was still apparent even at 0.4 µM LL-37. For the concentrations used, LL-37 primarily affects the initial or lag phase of growth.

**Figure 3.**
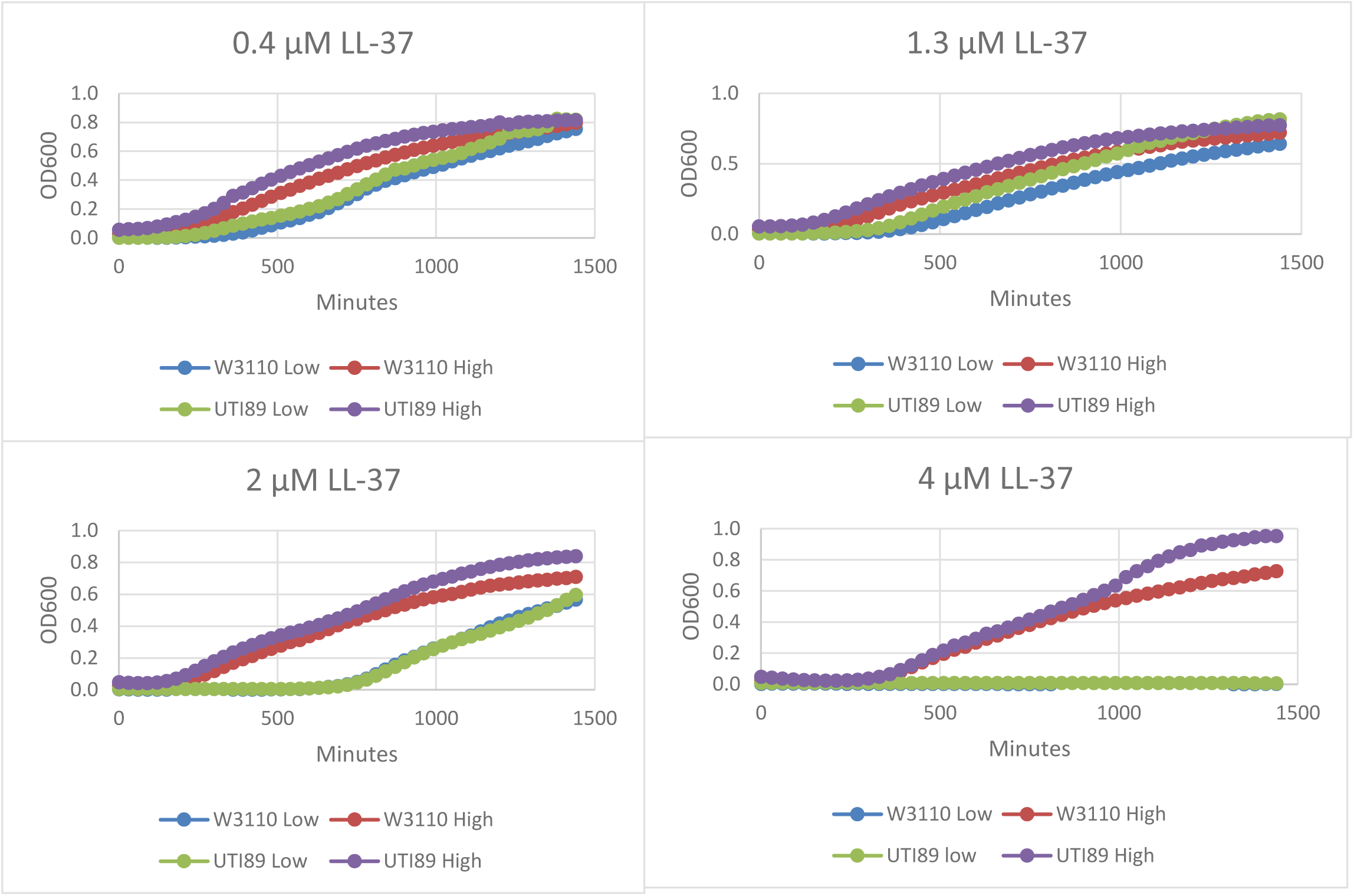
Effect of LL-37 on growth of UTI89 and W3110 in a minimal medium. The medium contained basic salts and glycerol as the carbon source. High and low refer to the inoculation density.

We analyzed growth in all urines after a low- and high-density inoculation as an indirect test for urinary IFs. We define low- and high-density inoculations as initial OD600s of 0.004 and 0.02, respectively, and the final growth ratio after a high versus low inoculation density as the HL ratio. For W3110, the HL ratios were high and variable which suggests sensitivity to IFs and substantial IF variation between individual urines (Table 3). For the UPEC strains, the average HL ratios were generally near 1.0 regardless of patient group (Table 3). Although the Figure 3 results are not consistent with the UPEC results, the in vitro results show an inoculation effect for LL-37 at an unphysiologically high concentration, while the low HL ratios for the UPEC strains grown in urine implies resistance to urinary IFs. These results do not imply that all UPEC strains are resistant to all IFs.

**Table 3.** The average HL ratio for each patient group. The indicated ratio is derived from the average HL ratio of 5 current and never group samples and 9 history group samples.

|  | W3110 | UTI89 | LRPF007 |
| --- | --- | --- | --- |
| Current | $3.9 \pm 4.0$ | $1.2 \pm 0.5$ | $1.2 \pm 0.3$ |
| History | $2.8 \pm 3.8$ | $1.3 \pm 1.0$ | $1.0 \pm 0.4$ |
| Never | $4.3 \pm 2.5$ | $1.0 \pm 0.1$ | $1.6 \pm 0.7$ |

### Urinary nutrient content between groups

If UPEC strains are resistant to IFs, then UPEC growth is a measure of utilizable nutrient content. The comparative growths of the two UPEC strains in current, never, and history group urines had ratios of approximately 3 to 2 to 1 (Table 4). For the current versus history groups, which has the highest ratio, the results achieve statistical significance for the high-density inoculations but not the low-density inoculations.

**Table 4.** Comparative growth in urine from different patient groups. The indicated ratios are derived from 5 current and never group samples and 9 history group samples. Because we are assessing nutrient content, the ΔOD600s were normalized to creatinine to adjust for hydration. F-tests were used to analyze if variance was unequal between the individual versus pooled urine samples. If the F-test was significant, the heteroscedastic *t*-test was used to test for differences in the means; if the F-test was not significant, the homoscedastic *t*-test was used. [note – no comparisons in this table needed the heteroscedastic *t*-test].

|  | UTI89 |  | LRPF007 |  | Average ratio |
| --- | --- | --- | --- | --- | --- |
| Inoculation density | Low | High | Low | High |  |
| Current vs History | 2.5 (0.090) | 3.0 (0.025) | 3.0 (0.15) | 2.9 (0.042) | 2.85 |
| Current vs Never | 1.4 (0.49) | 1.6 (0.35) | 1.5 (0.39) | 1.5 (0.73) | 1.5 |
| Never vs History | 1.7 (0.33) | 1.9 (0.19) | 1.8 (0.31) | 2.4 (0.14) | 1.95 |

In the nutrient-poor history group urine samples, both UPEC strains grew better than W3110 (Table 5). The UPEC strains also grew better than W3110 in diluted minimal and rich media (Figure 4), which shows that the UPEC strains utilize low levels of nutrients better than W3110.

**Table 5.**
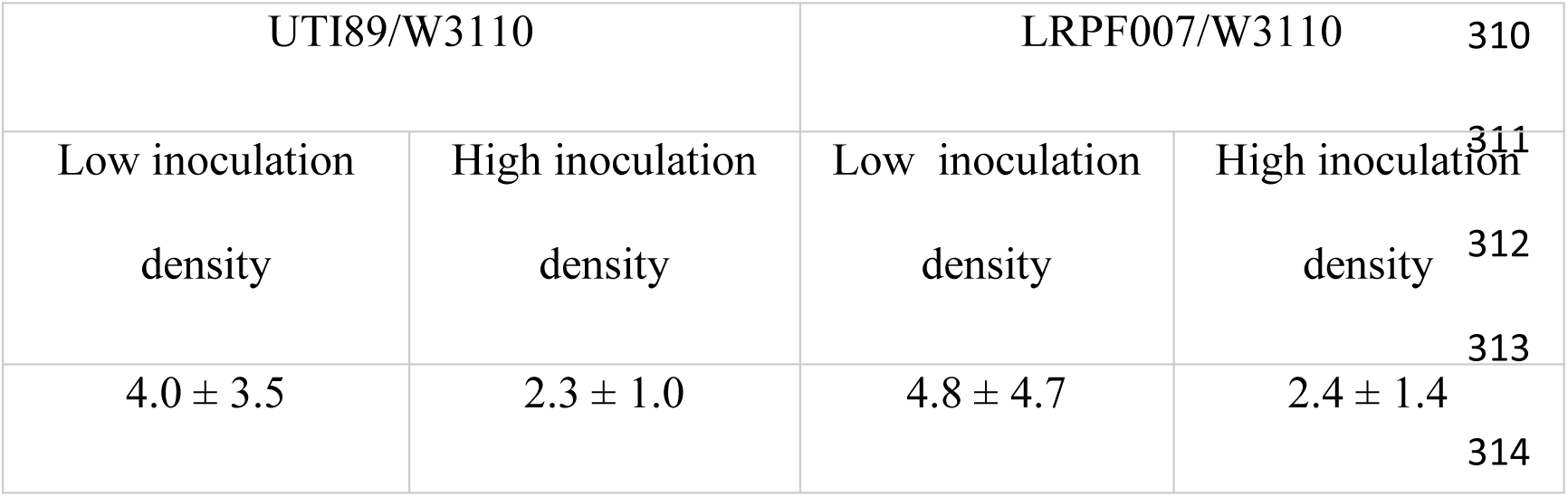
Comparative growth of the UPEC strains versus W3110 in history group urine. The ratios are based on 9 history group samples. The numbers are the average ratio ± SD.

| UTI89/W3110 |  | LRPF007/W3110 |  |
| --- | --- | --- | --- |
| Low inoculation<br>density | High inoculation<br>density | Low inoculation<br>density | High inoculation<br>density |
| $4.0 \pm 3.5$ | $2.3 \pm 1.0$ | $4.8 \pm 4.7$ | $2.4 \pm 1.4$ |

**Figure 4.**
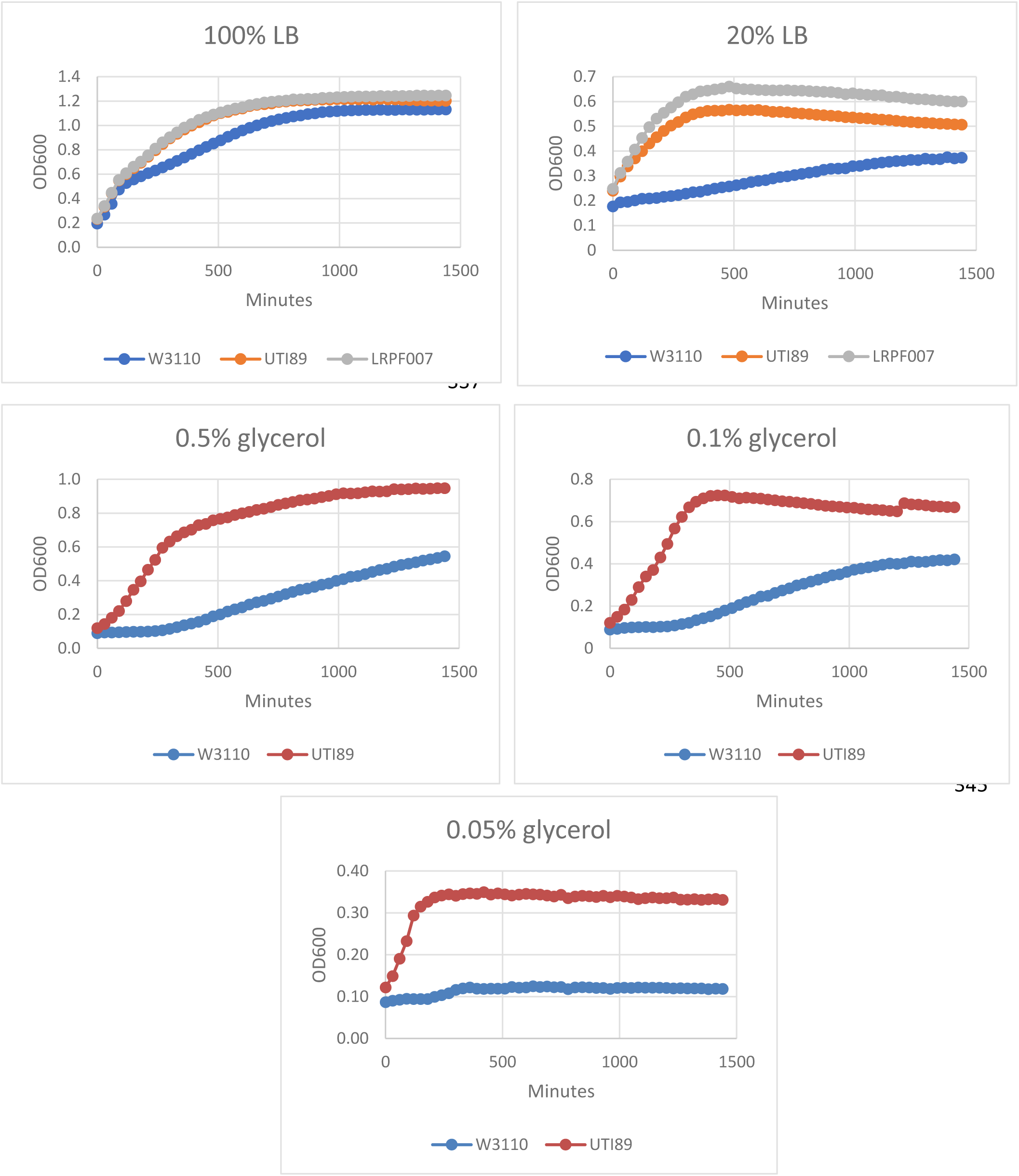
The growth of UTI89 and W3110 in diluted LB and minimal medium.

## DISCUSSION

We unexpectedly observed that bacterial growth in pooled urine is not necessarily the average growth of individual urines. In some cases, the pooled urine supported better growth than any individual urine. Of the two general factors that affect growth — IFs and nutrient content — only the latter can explain the growth synergism of the IF-insensitive UPEC strains. We propose that the pooling synergism results from optimization of a nutrient mixture. Current group urine samples do not show a pooling synergism, and we propose that pooling does not affect an already near optimal nutrient mixture. Perhaps current group urines contain an additional nutrient source such as lysed urothelial cells.

The amount of growth-promoting nutrients could explain the bacterial growth differences between the urine groups, especially between current and history group urine samples. Because of a small sample size, the ratio only sometimes reached statistical significance. If the high ratio reaches statistical significance with a larger sample size, then increased nutrient availability accompanies the transition from a dormant to an active infection.

The one study that analyzed nutrient composition and UTIs — a study of trauma patients who have a high incidence of UTIs — found elevated levels of several bacterial nutrients, including carbohydrates, amino acids, and iron (19). More knowledge of nutrient content could provide suggestions for interventions to prevent or better treat UTIs. Few studies have examined the relation between diet and UTIs (20, 21), but if nutrient content correlates with infection status, then understanding the relevant nutrients may suggest dietary changes that could affect UTI frequency.

As described above, the non-pathogenic W3110 is sensitive to the IFs present in our patient samples as assessed with the inoculation-density effect. W3110 growth might be useful for a rough estimate of total urinary IFs. We did not test whether W3110 is representative of other non-pathogenic strains, and our results provide no information about the IFs that are responsible for the W3110 inoculation effect. However, our observations suggest that W3110 growth might be useful for a rough estimate of total urinary IFs. A comprehensive study of urinary IFs has not been performed, although individual IFs have been studied. Two physiologically important antimicrobial peptides, ribonucleases 4 and 7, are higher in individuals who never have had a UTI than those who have a history of UTIs (22-24). Such results suggest that IFs may be an important inducible factor for UTI susceptibility as several studies indicate that an infection can increase IF levels (25-27). Further analysis is required to determine the primary urinary IFs, their variations between individuals, and their relevance to UTIs.

The growth synergism upon pooling has several important implications. First, it could defeat the purpose of pooling, if the objective is to obtain a representative urine. Pooled urine is often from healthy individuals (see Table 1), which could be in the never or history group, and pooling from these groups showed the greatest growth synergism. Second, considering the low number of participants in some studies that use pooled urine (Table 1), the implied assumption is limited variation in urine properties between individuals. However, our results showed large variations in urinary components between individuals within groups and between patient groups. These variations are consistent with a recent proposal that UTIs result from complex interactions between variable host and pathogen factors (10). Variable urinary composition is undoubtedly an important host factor. Third, interpretation of results from growth in pooled urine depends on whether pooled urine is an averaged urine or not. Expression of certain genes in a pooled urine may be a requirement for sustained growth, but such genes may not be expressed in an individual’s urine if growth is not optimal. Finally, reproducibility between labs is problematic if pooled urine is not a representative average urine. A mechanism to normalize results between labs might include an additional experiment with a synthetic urine which can be replicated in other labs.

Bacterial growth in urine depends on the urinary components and their concentration. Urine contains thousands of components which will vary between individuals and within one individual over time. Variation in urinary composition is, not surprisingly, one aspect of a complex host-pathogen interaction that affects the course of an infection. Our results are consistent with the possibility that changes in urinary components could trigger a latent infection or cause a relapse in patients with a history of UTIs. Further study is required to determine relevant urinary nutrients, and this knowledge could suggest non-antibiotic therapies to ameliorate UTI or minimize UTI recurrence.

## MATERIALS AND METHODS

### Urine collection and processing

Following IRB approval from both University of Texas Southwestern Medical School and University of Texas at Dallas, mid-stream urine samples were collected from post-menopausal women who had not been on antibiotics for at least 4 weeks. The urine was frozen at -80C. For the growth experiments, urine was thawed, centrifuged to remove particulates, and sterilized with a 0.2 µm filter. Each well in a microtiter dish received 0.2 ml urine.

### Bacterial strains

W3110 is a commonly used nonpathogenic lab strain. UTI89 is a highly passaged model UPEC strain (15). LRPF007 is a UPEC strain from a patient with RUTIs which was isolated after plating on ChromeAgar, streaked on L-broth to obtain single colonies, and cells from a single colony were grown in L-broth. Glycerol was added to the culture and the bacterial cells were frozen at -80 C. The strain was determined to be *E. coli* and phylotyped as described (28).

### Bacterial growth

Bacterial strains were streaked on an L-broth agar plate, grown overnight in L-broth medium, a single colony was inoculated into 1 ml L-broth for two hours in a 15 ml glass tubes at 250 rpm, centrifuged and resuspended in PBS three times to remove residual medium, resuspended in PBS, and then diluted either 100-fold or 20-fold into urine samples. Bacteria were grown in triplicate wells and optical density at 600 nm was measured in BioTek plate readers. The ΔOD600s ranged from 0.023 to 0.424. For cultures with a ΔOD600 > 0.1, the initial doubling time varied from 51 to 207 minutes and did not correlate with the ΔOD600. For the ΔOD600, the coefficients of variation (the ratio of the standard deviation to the average of the triplicate determinations) were low (Table 6).

**Table 6.** Coefficient of variation for growth in urine. The average coefficient of variation ± standard deviation.

| Growth | ΔOD600<br>range | Low-density inoculation<br>Average COV ± SD | High-density inoculation<br>Average COV ± SD |
| --- | --- | --- | --- |
| Bottom third | < 0.113 | 18 ± 15% | 18 ± 11% |
| Middle third | 0.117-0.236 | 13 ± 13% | 15 ± 19% |
| Top third | > 0.24 | 4.3 ± 1.8% | 9.5 ± 10.6% |

